# Magnetic wire active microrheology of human respiratory mucus

**DOI:** 10.1101/2021.04.05.438437

**Authors:** Milad Radiom, Romain Hénault, Salma Mani, Aline Grein Iankovski, Xavier Norel, Jean-François Berret

## Abstract

Mucus is a viscoelastic gel secreted by the pulmonary epithelium in the tracheobronchial region of the lungs. The coordinated beating of cilia in contact with the gel layer moves mucus upwards towards pharynx, removing inhaled pathogens and particles from the airways. The efficacy of this clearance mechanism depends primarily on the rheological properties of mucus. Here we use a magnetic wire based microrheology technique to study the viscoelastic properties of human mucus collected from human bronchus tubes. The response of wires between 5 and 80 µm in length to a magnetic rotating field is monitored by optical time-lapse microscopy and analyzed using constitutive equation models of rheology, including Maxwell and Kelvin-Voigt. The static shear viscosity and elastic modulus can be inferred from low frequency (10^−3^ − 10 rad s^−1^) measurements, leading to the evaluation of the mucin network relaxation time. This relaxation time is found to be widely distributed, from one to several hundred seconds. Mucus is identified as a viscoelastic liquid with an elastic modulus of 2.5 ± 0.5 Pa and a static viscosity of 100 ± 40 Pa s. Our work shows that beyond the established spatial variations in rheological properties due to microcavities, mucus exhibits secondary inhomogeneities associated with the relaxation time of the mucin network that may be important for its flow properties.

## 1 Introduction

The mammalian respiratory system is equipped with several defense mechanisms along the airways which serve to reduce the adverse effects of inhaled particulate matter. The first of these mechanisms is a physical barrier known as mucociliary clearance (MCC) in the tracheobronchial region.^1,2^ MCC consists of a recurring gel layer (mucus) secreted by the goblet cells and submucosal glands, and of metachronal waves of the cilia which expel the mucus from the airways. Mucus barrier has a thickness of about 2-5 μm, and is mainly composed of water and a network of mucin glycoproteins (5% wt. %).^1^ Much of the protective role of mucus is due to its adhesive properties as a result of high sugar concentration and a mesh structure.^3,4^ The efficacy of MCC depends primarily on the rheological properties of mucus.^5^ In this regard, several of pulmonary diseases such as asthma, cystic fibrosis and chronic obstructive pulmonary disease (COPD) are known to increase the mucus viscosity with secondary effects such as accumulation of pathogens and other inhaled particles leading to inflammation and epithelium damage.^6–8^

Rheological measurements of human pulmonary mucus are however difficult to carry out. One reason is the minute quantity of sample available from healthy individuals or diseased patients, typically of the order of a few microliters. Another reason is related to the mucus hydrogel structure, which contains nano-^9,10^ and microcavities^11,12^ in the range of 100 nm to 5 μm. These characteristics limit the application of both macro- and microrheology techniques to study the rheological properties of mucus.^1,5^ Motivated by the search for novel drug carriers for pulmonary therapies, bead tracking experiments, actually similar to those performed in passive microrheology, have been performed extensively in the recent years.^9–11,13–22^ These studies have demonstrated significant dependencies of the bead thermal diffusion on particle size and coating. It is now established that nanoparticles coated with poly(ethylene glycol) polymers can diffuse rather freely through the fluid containing microcavities of mucus. To overcome the difficulty related to rheology volume requirement, researchers have focused on sputum samples.^7,10,12,13,17,23–30^ The sputum samples were generally obtained from patients with serious lung conditions, such as cystic fibrosis and COPD. The collection methods were invasive and presented a risk of contamination with saliva or blood.^9,25^

We compiled sputum and mucus rheology data coming from 21 surveys.^9,10,12,13,15,23,25–39^ Data on cystic fibrosis sputum^7,12,13,23,27–30,32^ shows a gel-like behavior with the elastic modulus *G*′(*ω*) larger than the loss modulus *G*′′(*ω*) over the frequency range 0.01 – 100 rad s^−1^. At 1 rad s^−1^, it is found that the averaged values of *G*′ and *G*′′ are 4.9 ± 2.2 Pa and 3.6 ± 2.4 Pa, respectively, whereas the sputum shear viscosity is estimated to be 67 ± 34 Pa s at a shear rate of 1 s^−1^. The mucus data are in contrast less numerous and show viscosity values with larger uncertainties.^9,30,33,37,39^ The substantial extend of variations in the reported values is due to large inter-sample variations. This property of mucus makes interpretation of rheological properties difficult, especially for the design and development of mucolytic agents. From these investigations, both expectorated and native mucus have the characteristics of a viscoelastic gel.^40,41^

In this work, we investigate mucus rheological properties at length scales that are larger than the previously mentioned nano- and micro-cavities.^42,43^ We use microrheology platform called magnetic rotational spectroscopy (MRS) based on the time-lapse microscopy tracking of magnetically actuated micron-sized wires.^44–48^ *Ex Vivo* mucus samples are collected from human bronchus tube after surgery or from the culture of human bronchus tissue. The collected volumes in the former are in the milliliter range, while the volumes secreted in the latter case are in the range of 10 – 30 μl, which is sufficient for MRS. Wires of length 5 – 80 μm, *i.e.* longer than the typical mucus microcavities, and diameter 0.7 – 2 μm are added to mucus and submitted to an external magnetic field of increasing frequency. The wire response is monitored by time-lapse optical microscopy and analyzed using constitutive equation models of rheology, including Maxwell and Kelvin-Voigt.^44,49,50^ In particular, with measurements made at frequencies of the order of 10^−3^ rad s^−1^, the static shear viscosity and elastic modulus can be inferred,^49,50^ leading to the evaluation of the mucin network relaxation time. In what follows, we present the results obtained with mucus and discuss the presence of viscoelastic heterogeneities in terms of relaxation times.

## 2 Materials and Methods

### 2.1 Mucus collection

Early and late *Ex Vivo* mucus were collected as follows. A human lung was received after transplantation surgery from the anatomic pathology service of Hospital Bichat-Claude-Bernard in Paris, France. The bronchus tube of the lung was initially inspected to be in a healthy condition. The mucus that was secreted inside the tube was gently taken using tweezers – this mucus is called early *Ex Vivo*. To prepare late *Ex Vivo* mucus, a part of a bronchus was then excised to tubes of length 7.5 – 10 mm and diameter 5 – 10 mm. These tubes were further cut into several smaller rings as shown in Fig. 1. The rings were immersed in Tyrode’s solution and incubated for 18 h inside a CO_2_ incubator at 37 °C. During this incubation period, the epithelial mucous (goblet) cells continue to produce mucus. The secreted mucus whose volume was in the range of 10 – 30 μl was gently collected using tweezers and kept at 4 °C until microrheology experiments within 24 h after collection (**Supporting Information S1**). This procedure was applied to lungs received from 8 different patients who underwent operations between January and April 2019). As the microrheology results between the patients did not reveal any notable differences, the data of the different patients were pooled together and treated as a unique batch. The above procedures are approved by the Institutional Review Board of the Hôpitaux Universitaires Paris Nord, Université de Paris and Assistance Publique - Hôpitaux de Paris (AP-HP). Prior to surgery, the patients signed a written informed consent to participate in scientific studies (**Supporting Information S2**).

**Figure 1:**
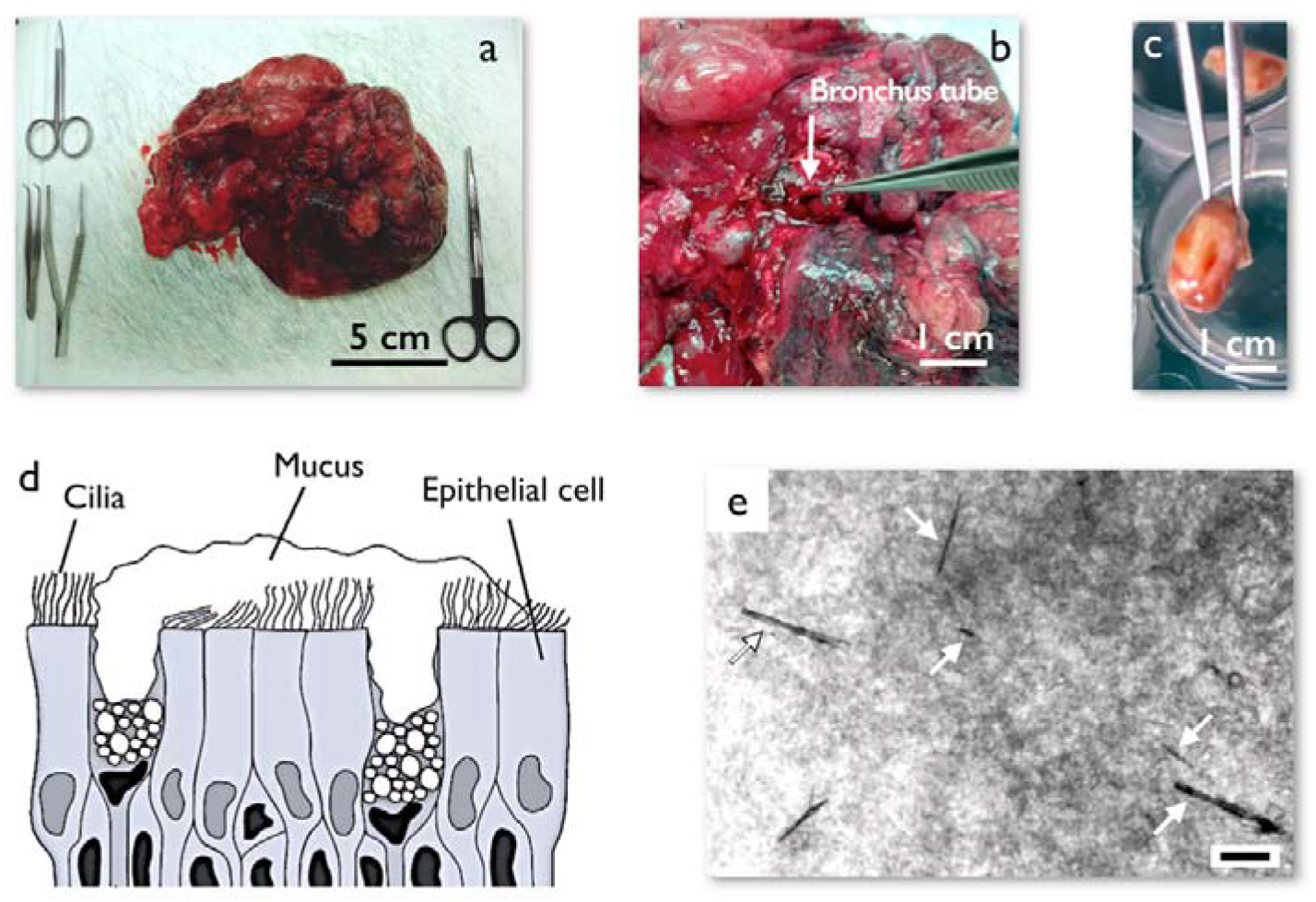
a) Piece of a human lung after surgery obtained from the anatomical pathology service at Hospital Bichat-Claude-Bernard, Paris, France. b) Dissection of a human lung showing a bronchus tube. c) A cut section of human bronchus tube from a transplanted lung after surgery. This section is incubated in Tyrode’s solution for 18 h during which time it secrets 10 – 30 μl of mucus. d) Schematic of the epithelium including mucus secreting goblet cells, ciliated cells and mucus lining. Mucus contains mucin glycoproteins which form a porous network with pore sizes in the range of 100 – 5000 nm^11^. (e) Phase contrast microscope image of mucus with injected micron-sized magnetic wires. The wires are marked with arrows. The mucus has a heterogeneous structure that is observable under phase contrast microscopy. The scale is 10 μm.

### 2.2 Mucus dry mass

When the mucus samples were in sufficient quantity (> 70 μL) the dry mass was also quantified. For this purpose, the remaining mucus from sample preparation for magnetic rotational spectroscopy (MRS) was heated to 150 °C for 1 h and then weighed to obtain the ratio of the dry mass to the total mass. Experiments were performed in triplicate on the *early Ex Vivo* mucus, leading to mass ratios equal to 4.3 %, 5.7% and 5.3% (average 5.1 ± 0.6%), in good agreement with literature.^1^

### 2.3 Sample preparation for magnetic rotational spectroscopy experiments

All sample preparations were performed inside a laminar flow hood in a cell culture room. The vials containing the mucus were shut at all time and only opened when wires were added to mucus and prior to deposition on glass coverslip. This step was necessary to minimize the evaporation of the water content of mucus. We adhered a dual adhesive Gene Frame (thickness 250 μm, Abgene, Advanced Biotech) to a cover slip. Afterwards, 0.5 μl of a solution containing magnetic wires was added to 10 – 30 μl of mucus and the sample was deposited in the center of the Gene Frame. The sample was then covered by a second coverslip and hermetically sealed.

### 2.4 Synthesis of magnetic wires

Superparamagnetic maghemite (*γ*-Fe_2_O_3_) nanoparticles with a mean diameter of 13.2 nm and a dispersity of 0.23 were used. The nanoparticles were initially coated with poly(acrylic) acid polymers^51,52^ (Aldrich, France) of molecular weight 2100 g mol^−1^. The negatively charged particles were then assembled using poly(diallydimethyl ammonium chloride) (PDADMAC, *M*_*w*_ > 100000 g mol^−1^, Aldrich, France). The cylindrical shape was given during desalting the mixed solutions using a Slide-a-Lyzer^®^ dialysis cassette (Thermo Fisher Scientific, France) with molecular weight cut-off equal to 10000 g mol^−1^ under a magnetic field of 0.3 T (**Supporting Information S3**).^51^

### 2.5 Characterization of magnetic wires

The geometrical characterization of the wires was performed by measuring the length *L* and the diameter *D* of wires using a 100× objective lens on an optical microscope (Olympus IX73) coupled with a CCD camera (QImaging, EXi Blue) supported by the software Metaview (Universal Imaging). The 100× objective with numerical aperture 1.3 gives an (X,Y)-spatial resolution of 280 nm. From these measurements, we calculated the reduced wire length, 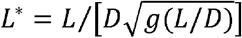 and plotted it as a function of the actual length, leading to a scaling of the form *L*\* = 1.66*L*^0.64^. The anisotropy of magnetic susceptibility Δ*χ* featuring in the expression of the critical frequency *ω*_*C*_ (Eq. 1) and of the asymptotic return angle *θ*_*B*_ (Eq. 2) was calculated in independent measurements in fluids of known viscosity, such as water-glycerol mixtures.^44^ By collecting *ω*_*C*_ for wires of various *L*\*, Δ*χ* was calculated using Eq. 2 in the main text. For these wires Δ*χ*. was found to be equal to 2.3 ± 0.5 (**Supporting Information S4&S5**).

### 2.6 Wire rotation measurements on mucus

The magnetic wires used in this study have lengths between 5 and 80 μm and diameters between 0.7 and 2 μm. The cell containing the mucus sample with embedded wires was introduced into a homemade device generating a rotational magnetic field, from two pairs of coils (resistance 23 Ω) working with a 90°-phase shift. An electronic set-up allowed measurements in the frequency range *ω* = 10^−3^ − 10^2^ rad s^−1^ and at a magnetic field *μ*_0_*H* = 10.3 mT (**Supporting Information S6**). The wire rotation was monitored in phase contrast mode with a 20× objective. Because the (X,Y)-spatial resolution of this objective (920 nm) is insufficient to measure the diameter of each wire with accuracy, the scaling expression *L*\*(*L*) determined with the 100× was used for the data treatment. For each angular frequency, a movie was recorded for a period of time of at least 10/*ω* and then treated using the ImageJ software to retrieve the wire rotation angle *θ*(*ω, t*). The above experimental procedure was applied to a total of 98 wires, 58(resp. 40) in early (resp. late) *Ex Vivo* mucus, each of which was studied at 4 - 5 different angular frequencies.

### 2.7 Comparison of magnetic rotational spectroscopy with cone-and-plate rheology

Wire rotation may be interpreted in terms of several models such as pure viscous (Newton), viscoelastic (Maxwell, Kelvin-Voigt) and pure elastic (Hooke) responses as shown in **Supporting Information S7**. The validity of approach and data analysis was confirmed in experiments performed in wormlike micellar solutions and polysaccharide gels as models of viscoelastic liquid and viscoelastic solid which are detailed in **Supporting Information S7.** As a result, it was shown that the MRS technique measures the linear rheology response parameters,^40^ such as the static shear viscosity and the instantaneous and equilibrium elastic shear moduli, *G*_0_ and *G*_*eq*_ respectively.

## 3 Results and Discussion

### 3.1 Mucus collection and wire seeding

Mucus samples were obtained from the anatomical pathology service at Hospital Bichat-Claude-Bernard in Paris. The samples were taken in two manners: one directly from bronchus tubes after surgery and the other from the culture of bronchus tissue rings. The first mucus is called early *Ex Vivo* mucus and the second one, late *Ex Vivo* mucus. The collected lung was dissected to extract pieces of bronchus tubes which were then cut into rings of length 2 – 5 mm. The rings were cultured for 18 h in Tyrode’s solution, during which time they produced their own mucus (Figs. 1a-1c). The collection procedures and conditionings of the different samples are summarized in **Supporting Information S1**. A schematic of the cultured tissue is depicted in Fig. 1d. It is composed of ciliated cells as well as mucus secreting goblet cells. The readily polarized goblet cells of the cultured tissue produce mucus towards the lumen side. After collection, a small volume of wire containing solution (0.5 μl) was then added to the sample. An example of wire distribution in mucus is shown in Fig. 1e. We observed that the wires spread somewhat uniformly in the sample. Additionally, wires could be found over the full thickness of mucus sample; however, the measurements were performed on wires located in the center of the sample to reduce wall interaction effects. Occasionally we found large number of wires co-located in one region of mucus. These wires were not selected for microrheology investigation due to possible hydrodynamic interactions. Fig. 1e further shows that the background mucus interacts differently with passing light in local regions (*i.e.* the background is not uniform). This observation manifests that mucus has a heterogeneous material distribution, most likely related to the microcavities mentioned in the introduction.

### 3.2 The different regimes of wire rotation in human respiratory mucus

In this study, we investigated the rotation of a total of 98 magnetic wires dispersed in human mucus samples, 40 from the late *Ex Vivo* and 58 from the early *Ex Vivo* mucus. Each wire was studied at 4 to 5 frequency between 2 10^−3^ and 20 rad s^−1^. For these 98 wires, we found two generic rheological behaviors, namely that of viscoelastic liquids^49^ and that of soft solids.^50^ Interestingly, none of the wire studied showed rotation features of a purely viscous liquid.^53^ Magnetic wires indeed exhibit specific angular frequency dependencies whether they are embedded in one or the other of the materials mentioned above. For viscoelastic liquids, at low angular frequency *ω*, the wires rotate synchronously with the field, and above a critical frequency *ω*_*C*_ they show asynchronous motion with back-and-forth oscillations. In that case, the average rotational velocity, defined as Ω(*ω*) = < *dθ*(*ω, t*)/*dt* >_*t*_, where *θ*(*ω, t*) is the orientation angle of the wire, is different from zero at all values of (l). For soft solids, a unique response is found as a function of the frequency, namely that of an asynchronous oscillation associated with Ω(*ω, t*) = 0. This specificity of soft solids by MRS comes from the fact that at the magnetic fields used, the stresses applied locally by the wires are lower than the yield stress and that we are dealing with in a purely deformational regime, and not with a flow regime.^50^ Of note, each mucus sample studied exhibited both rheological behaviors, an outcome which will be explained later on the basis of a quantitative analysis of the *θ*(*ω, t*)-times series. Table I shows the number of wires entering each rheological class for late and early *Ex Vivo* mucus. We have found that samples taken from lung after surgery were contaminated with blood and possibly other lung fluids (**Supporting Information S1**). It is hypothesized that its rheological property may be impaired and do not reflect that of intact mucus. For this reason, we focus first on the late *Ex Vivo* mucus cultured from bronchus tissues.

**Table I:** Summary of the number of wires exhibiting viscoelastic liquid and soft solid rheological behaviors.

| Mucus type | Number of wires with viscoelastic liquid behavior | Number of wires with soft solid behavior | Total |
| --- | --- | --- | --- |
| Late *Ex Vivo* mucus | 24 | 16 | 40 |
| Early *Ex Vivo* mucus | 35 | 23 | 58 |

**Table II:** Combination of the rheological parameters obtained for late and early Ex Vivo mucus using the magnetic rotational spectrometry technique. Here, η, G and τ denote the static shear viscosity, the elastic modulus and the relaxation time, respectively.

| Mucus type | Viscoelastic liquid behavior |  |  | Soft solid behavior |
| --- | --- | --- | --- | --- |
| | $\eta$ (Pa s) | $G$ (Pa) | $\tau$ (s) | $G$ (Pa) |
| Late *Ex Vivo* mucus | $100 \pm 40$ | $2.5 \pm 0.5$ | $47 \pm 12$ | $2.6 \pm 0.3$ |
| Early *Ex Vivo* mucus | $180 \pm 30$ | $4.4 \pm 0.5$ | $49 \pm 6$ | $2.0 \pm 0.2$ |

### 3.3 Late *Ex Vivo* human mucus with viscoelastic liquid behavior

#### Viscosity measurements

We will initially describe the features corresponding to the viscoelastic liquid behavior. Fig. 2a illustrates a 23 μm wire under a 10.3 mT rotating magnetic field and frequency equal to 0.0094 rad s-1. One observes a heterogeneous background under phase contrast microscopy which is associated with a non-uniform material distribution. It is also noted that the sizes of the heterogeneous regions (~ 1 μm) are smaller than the length of the wire. The different images of the chronophotograph at time intervals of 100 s show that the wire rotates synchronously with the field. By increasing the frequency to 0.94 rad s^−1^ (Fig. 2b), the wire exhibits a transition to asynchronous rotation whereby it performs back and forth oscillations: after an initial increase in the orientation angle in the direction of the field (yellow arrow), the wire undergoes a subsequent return in the opposite direction (red arrow) followed by a next rise in the orientation angle. We refer to the transition from the synchronous to the asynchronous regime by a frequency that we denote as critical frequency, *ω*_*C*_. For the wire in Fig. 2, *ω*_*C*_ is estimated to be 0.12 rad s^−1^.

**Figure 2:**
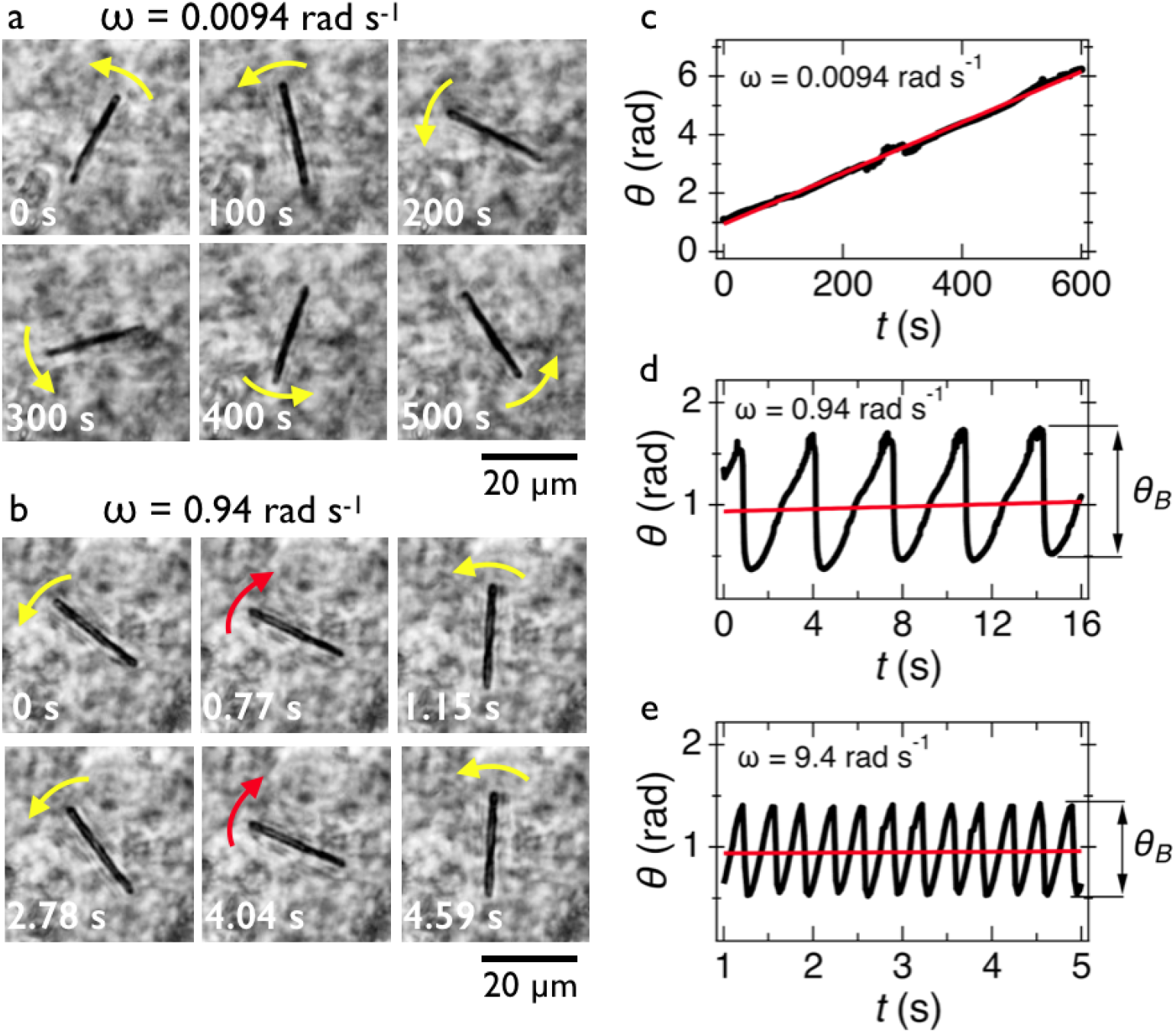
**a-b)** Chronographs of a 23 μm wire undergoing rotation in Ex Vivo mucus as a result of a 10.3 mT external magnetic field rotating with a frequency of 0.0094 rad s^−1^ (a) and 0.94 rad s^−1^ (b). **c-e)** Time series of orientation angle in the synchronous (c) and asynchronous regimes (d and e). The average rotational velocity is depicted via straight red lines and the oscillation amplitude by arrows.

In Fig. 2c, we show the *θ*(*t*)-time series at *ω* = 0.0094 rad s^−1^ below the critical frequency. The orientation angle rises linearly with time and the slope of the temporal evolution is shown by a straight red line denoting *Ω*. For *ω* < *ω*_*C*_, we find *Ω*(*ω*) = *ω*. In Figs. 2d and 2e, we show the *θ*(*t*)-time series at *ω* = 0.94 rad s^−1^ and 9.4 rad s^−1^, whereby *Ω* is now found to be 0.015 rad s^−1^ and 0.006 rad s^−1^, respectively (red lines). Above the critical frequency (*ω* > *ω*_*C*_), *Ω* becomes smaller than *ω* and follows the relation^47,53^ 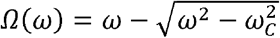. In the asynchronous regime, the oscillation amplitude *θ*_*B*_ as depicted by arrows is the angle by which the wire turns in the opposite direction to the field, here about 1.2 rad and 0.9 rad, respectively. The movies associated with Figs. 2a and 2b are provided in **Supporting Movies 1 & 2**.

Figs. 3a, 3b and 3c show the evolution of for wires with lengths 10, 24 and 33 μm, respectively. One observes that increases linearly with the actuation frequency in the synchronous regime ( ) until it passes through a cusp-like maximum at the critical frequency and then falls rapidly ( ). This behavior is used to interpret the critical frequency for each wire. Fig. 3d shows the normalized velocity as a function of normalized frequency for the 24 wires which share this behavior. One observes that the normalized wire responses agree reasonably with each other, especially considering the heterogeneity of mucus and its non-uniform material distribution. Increased dispersity at high frequencies is associated with a lower precision in the determination of *Ω* from wire tracking microscopy in this part of the spectrum.

**Figure 3:**
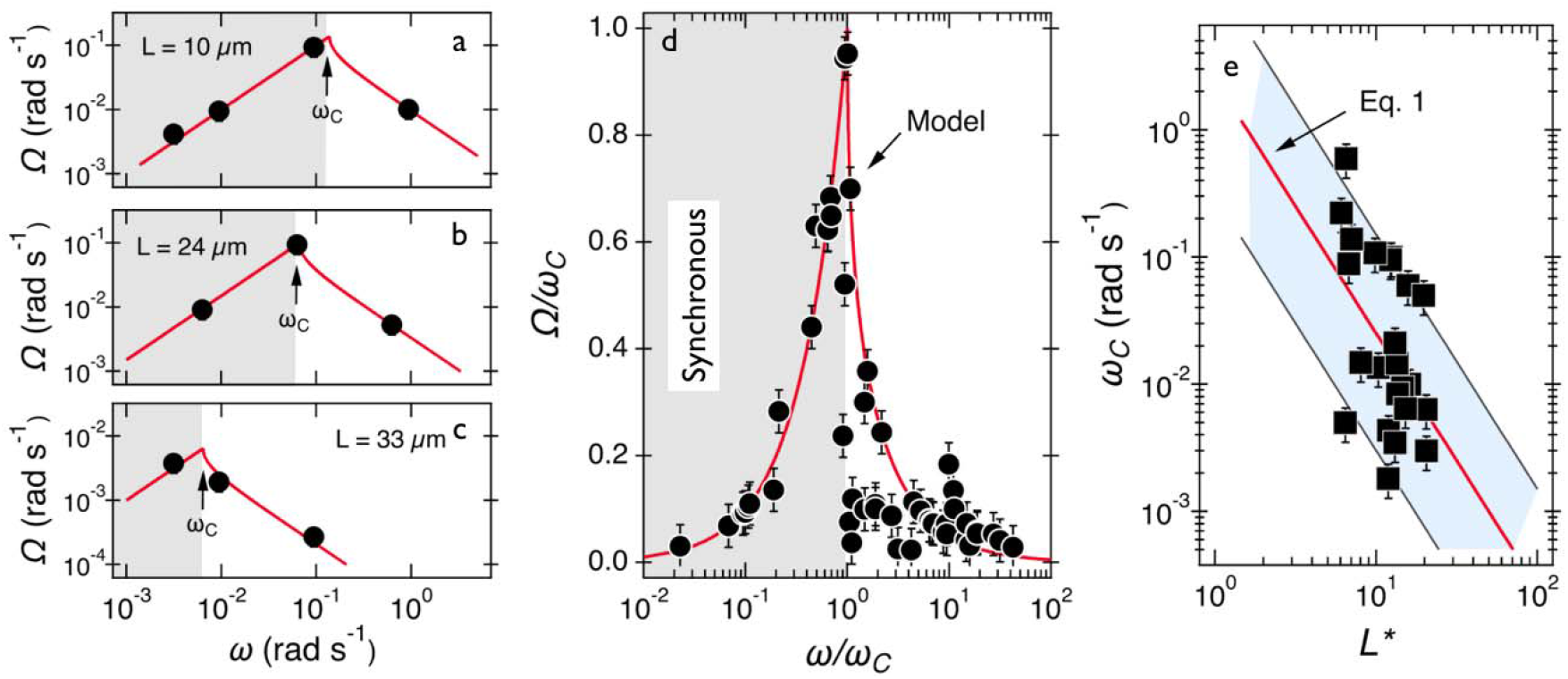
**a-c)** Average rotational velocity of several wires with length between 10 and 33 μm. Solid fit lines are from the expressions given in the text. **d)** Average rotational velocity of wires normalized by the critical frequency plotted versus normalized frequency. **e)** Variation of critical frequency as a function of reduced wire length. The thick solid line shows the -dependence corresponding to a mean viscosity = 100 ± 40 Pa s.

The viscosity of mucus can be identified from the asymptotic behavior of *ω*_*C*_ as a function of the wire length. As shown previously^49^ *ω*_*C*_ is related with the static viscosity *η via* the expression:

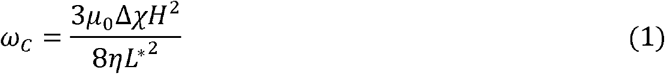

where 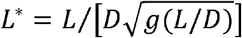 is the reduced wire length, *g*(*x*) = *ln*(*x*) − 0.662 + 0.917*x* − 0.050*x*^2^, and *L* and *D* the wire length and diameter, *μ*_0_ the vacuum permeability, *H* the magnetic field strength and Δ*χ* the anisotropy of magnetic susceptibility. Fig. 3e displays *ω*_*C*_ as a function of *L*\* for the same wires shown in Fig. 3d. In this figure, each data point represents a single wire embedded in a different mucus environment. Using Eq. 1 we calculated the static viscosity of the individual wires and obtained a mean value equal to 100 ± 40 Pa s (± error of mean, 6 – 280 Pa s are the limits of the shaded area in Fig. 3e). The viscosity that we measure is much larger than those reported from particle tracking (0.05 – 0.l Pa s) and also slightly larger than that of sputum (67 ± 34 Pa s) obtained from macrorheology (see Section *Comparison with literature data*). This outcome signifies the success of the MRS technique in measuring the viscosity of a heterogeneous sample from the local rotation-actuation response of magnetic wires.

#### Elastic modulus measurements

Figs. 4a, 4b and 4c show as a function of for wires with lengths 12, 27 and 52 μm, respectively. As a general behavior, decreases with frequency and we denote the value at the highest actuation frequency by. Fig. 4d displays as a function of the normalized frequency. It is found that the data superimpose over 4 decades in frequency and follow a scaling law with exponent −0.10. On the same figure we also show the solution of the constitutive equation for Newton fluids, which predicts a fast fall of following the relation for. The variation of the oscillation amplitude as observed with wires in this regime is a signature of a viscoelastic liquid.^44,49^ The elastic modulus can be interpreted from the asymptotic behavior of as a function of the wire length. As shown previously,^50^ this angle is related with the elastic modulus *via* the expression:

**Figure 4:**
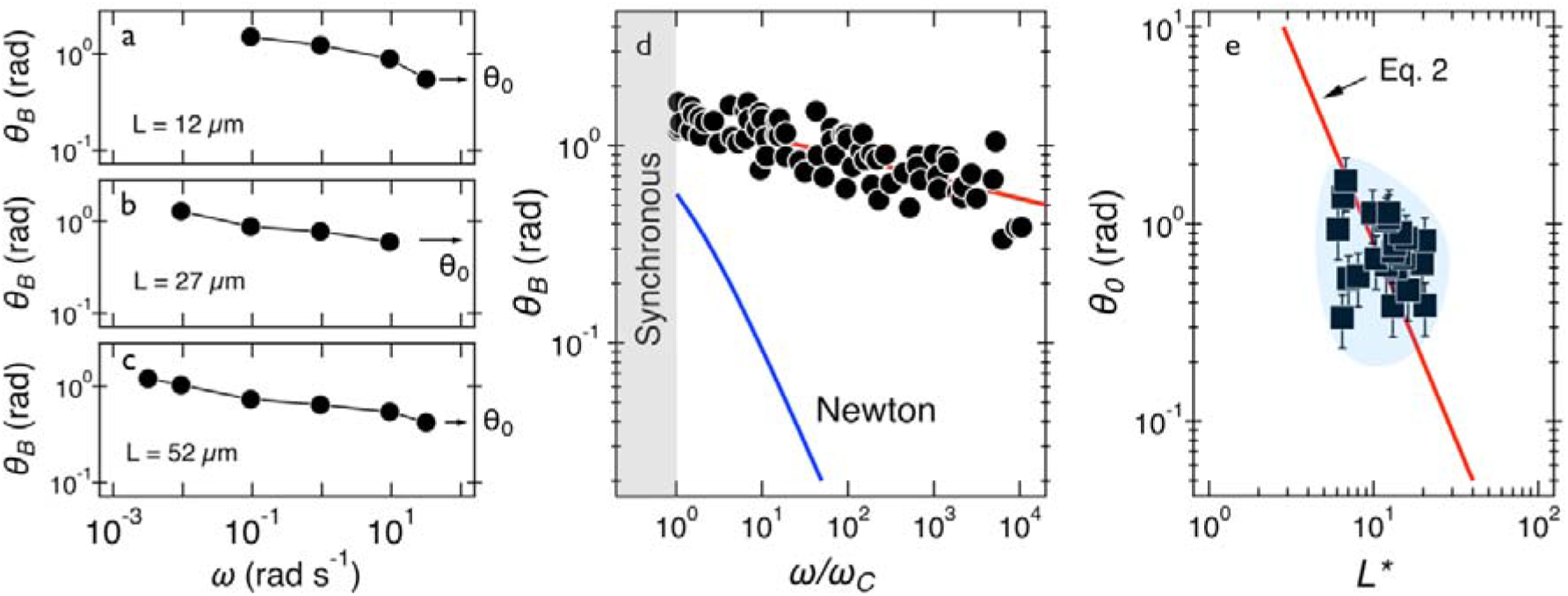
**a-c)** Oscillation amplitude of several wires with lengths 12 – 52 μm. Arrows point at the oscillation amplitude at the highest frequency, denoted by. **d)** Oscillation amplitude as a function of normalized frequency and comparison that of a Newton fluid. **e)** Variation of as a function of reduced wire length. Solid line shows the - dependence of Eq. 2, leading to = 2.5 ± 0.5 Pa. In this figure, each data point represents a different wire in mucus.

Fig. 4e displays this angle for the 24 individual wires of this class as a function of the reduced wire length and also depicts the -dependence of this angle. The shaded area in this figure shows the extent of variation of the prefactor in Eq. 2. Using Eq. 2, we calculated the elastic modulus and obtained a mean of = 2.5 ± 0.5 Pa (0.5 – 13 Pa are the limits of the shaded area). The variation in elasticity values similar to the viscosity is due to mucus heterogeneity at the scale of the wire length, *i.e.* in the range 5 – 80 μm.

### 3.4 Late *Ex Vivo* human mucus exhibiting soft solid characteristics

Out of 40 wires investigated in late *Ex Vivo* mucus, 16 wires with length between 8 and 75 μm showed a soft solid behavior characterized by an average rotational velocity close to zero, or precisely = 10^−4^ rad s^−1^, at all values of. This limitation is obtained from the resolution of the optical microscope and that of the wire orientation obtained from the tracking ImageJ plugin.^54^ During observation periods which can be longer than 1 h the wire remained stationary and only performed an oscillatory motion. Figs. 5a and 5b show examples of such behavior for a wire with length 23 μm under actuation fields with *ω* = 0.0031 rad s^−1^ and 0.0094 rad s^−1^ respectively. These observations are generally indications that a viscoelastic (soft) solid material surrounds the wire. From this type of response, a critical frequency cannot be assigned. It is possible however to obtain *θ*_*B*_(*ω*) and calculate *G* from Eq. 2. Fig. 5c displays *θ*_*B*_(*ω*) for wires with lengths between 14 μm and 75 μm and Fig. 5d shows the *θ*_0_(*L*\*)-dependence that corroborates the scaling of Eq. 2.^50^ We calculated *G* for individual wires and obtained a mean of *G* = 2.6 ± 0.3 Pa. A similar calculation was made for the low frequency region, from which the equilibrium modulus can be derived^44,50^ *via* the relation lim_*ω*→0_*θ*_*B*_(*ω*) = 3*μ*_0_Δ*χH*^2^/4*L*\*^2^*G*_*eq*_. It is found that *G*_*eq*_ = 1.7 ± 0.3 Pa, *i.e.* a value close to that of *G*. The movies associated with the wire rotations in Figs. 5a and 5b are shown in **Supporting Movies 3 & 4**.

**Figure 5:**
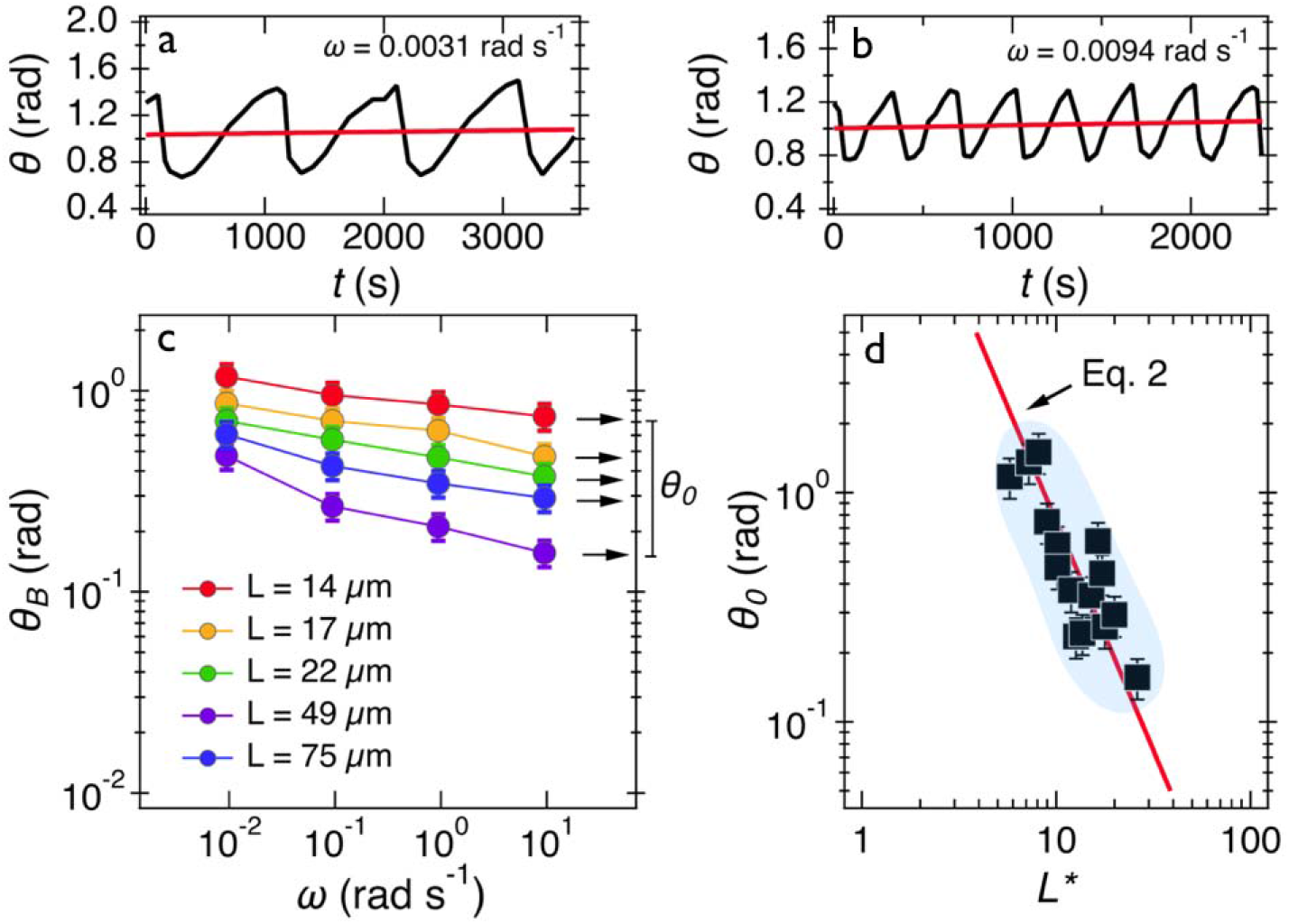
**a, b)** Time dependence of orientation angle for a wire experiencing the second generic behavior at actuation frequencies 0.0031 rad s^−1^ and 0.0094 rad s^−1^ respectively. **c)** Oscillation amplitude of wires with lengths 14 – 75 μm. Arrows point at the amplitude at the highest frequency. **d)** Variation of oscillation amplitude as a function of the reduced wire length. Solid line shows the -dependence of Eq. 2, leading to = 2.6 ± 0.3 Pa. In this figure, each data point represents a different wire dispersed in mucus.

We thereby find that the elastic moduli measured at regions with local viscoelastic liquid or soft solid behavior are in good agreement. This agreement shows that the elastic networks of these mucus regions with these unlike behaviors are similar. We therefore assume that these differences come from the relaxation times. For the viscoelastic fluid case, the relaxation time of mucus can be estimated from the values of the viscosity and the elastic modulus, *τ* = *η*/*G* for each wire, leading to a mean relaxation time equal to *τ* = 47 ± 12 s (min = 1 s, max = 290 s). For the soft solid case, we can assign an upper limit for *ω*_*C*_, of the order of the minimum angular frequency used *i.e.* 10^−3^ rad s^−1^. This outcome indicates that the current MRS technique is limited to measurements of relaxation times less than 300 s. To probe such long relaxation times, a steady rotation of wires could be achieved if the wire length would be reduced but this has the disadvantage of approaching the size of microcavities, or if the angular frequency would be decrease below 10^−3^ rad s^−1^.

### 3.5 *Ex Vivo* human mucus collected after surgery

We also investigated mucus that was directly taken from the bronchus tube after surgery, *i.e.* the early *Ex Vivo* mucus. Similarly, we find two generic behaviors from wire rotations. Out of the 58 wires investigated in 8 samples, 35 presented the response of a viscoelastic liquid and 23 showed the response of a soft solid. Note, the proportion between the two types of responses was similar that of the late *Ex Vivo* mucus, around 60% (Table I). In the first generic behavior (*i.e*. that of the viscoelastic liquid), the static viscosity was calculated to be = 180 ± 30 Pa s from Eq. 1 and the elastic modulus to be = 4.4 ± 0.5 Pa from Eq. 2. In the second generic behavior (*i.e.* that of the soft solid) we calculated an elastic modulus equal to *G* = 2.0 ± 0.2 Pa. Similarly, we assign a relaxation time to the first generic behavior equal to *τ* = 49 ± 6 s and an upper limit for the relaxation time in the second generic behavior estimated to be > 200 s. Details of the experiments and results with early *Ex Vivo* mucus are shown in **Supporting Information S8**. We find that the viscoelastic properties of early *Ex Vivo* mucus agree reasonably with those calculated in late *Ex Vivo* mucus, indicating that the contamination identified on samples taken after surgery have little to no effect on the rheology (**Supporting Information S1**). In addition, the late *Ex Vivo* mucus is similar in production to cell culture methods certifying the applicability of our protocol in investigations related to mucus flow in airway surfaces.

### 3.6 Comparison with literature data

Following this study, it was important to compare our *Ex Vivo* mucus microrheology data on to that of the literature. To this end, macro and microrheological data from 21 sputum and mucus surveys were evaluated.^9,10,12,13,15,23,25–39^ The nature, origin and references of the samples selected for this analysis are listed in **Supporting Information S9**. From these reports, the elastic modulus *G*′(*ω*), the loss modulus *G*′′(*ω*) and the complex modulus were compiled. Whenever this was possible, the angular frequency dependence was taken into account and the data was extrapolated to = 1 rad s^−1^ to allow comparison. For cystic fibrosis sputum (for which the number of samples were the most numerous), averaged moduli, and were obtained from = 7, 5 and 8 independent studies respectively. It was found that at 1 rad s^−1^, = 4.9 ± 2.2 Pa, = 3.6 ± 2.4 Pa and = 12.4 ± 4.3 Pa. For mucus samples, the following results were obtained: = 7.3 ± 2.9 Pa, = 2.1 ± 0.7 Pa and = 10.7 ± 2.6 Pa from = 3, 3 and 5 independent studies respectively. Finally, the shear viscosity obtained at low shear rate (< 0.1 s^−1^) was also evaluated and found to be 67 ± 34 Pa s for cystic fibrosis sputum (= 5) and 382 ± 286 Pa s for mucus (= 4). Figs. 6a and 6b show the moduli histograms (in dark blue, in green and in orange) for the cystic fibrosis sputum and mucus, whereas Figs. 6c compares the shear viscosity of these two lung fluids. Given the uncertainties from the literature data, our results (in grey in Figs. 6c) are in overall agreement with those reported in earlier studies, demonstrating that the MRS method is reliable and sensitive.

**Figure 6:**
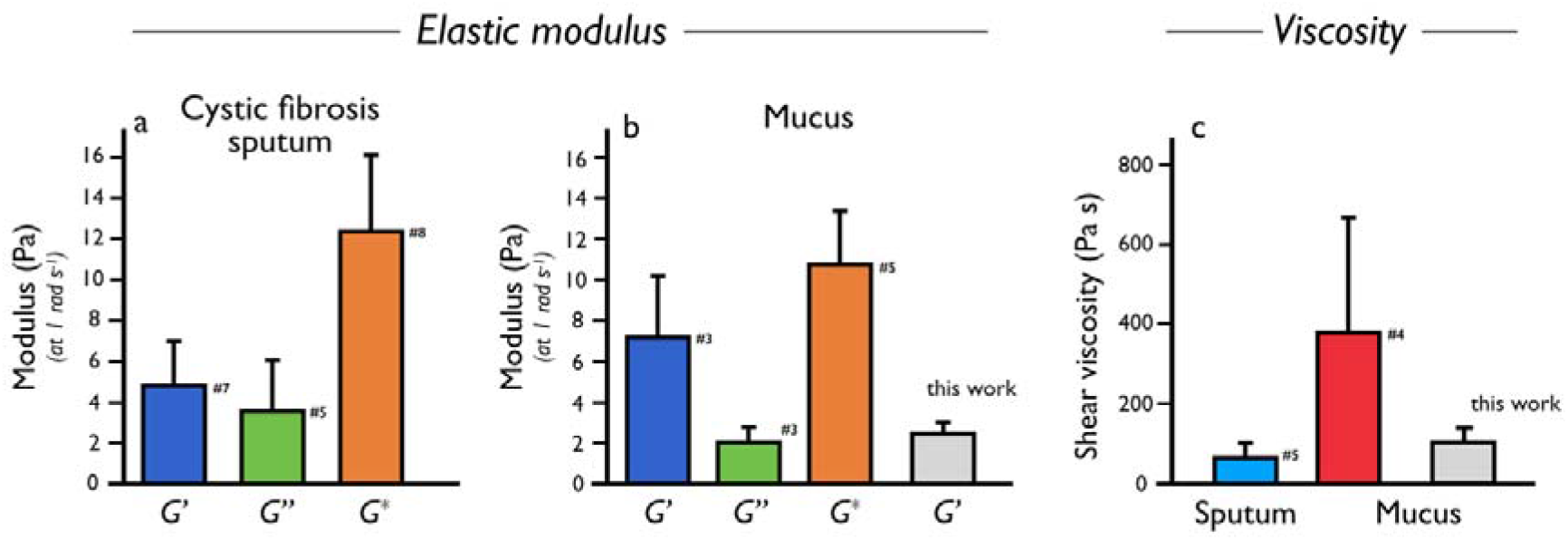
This figure is for human sputum and mucus and corresponds to the data collected from **Table S9**. The elastic ( , dark blue), loss ( , green) and complex ( , orange) moduli for cystic fibrosis sputum (a) and mucus (b). Comparison between the shear viscosity reported in the literature for sputum and for mucus (c). Note the large values of the standard errors for mucus shear viscosity.

## 4 Conclusion

In this work we propose a new active microrheology technique to determine the rheological characteristics of human respiratory mucus. The probes used are remotely actuated magnetic wires of lengths between 5 and 80 μm, *i.e.* much greater than the size of the microcavities (100 nm - 5 μm) commonly observed in mucus samples.^11^ We thereby ensure that the viscosity and elasticity values obtained are averaged over length scales large enough to be compared with the bulk values. One result that emerges from this work is that human respiratory mucus is a viscoelastic material, an outcome that corroborates earlier literature reports.^5^ The ability to perform MRS experiments at angular frequency as low as 10^−3^ rad s^−1^ makes it possible to highlight the viscoelastic liquid character of the mucus, and also to estimate for the first time the relaxation times of the mucin network. In human respiratory mucus, we have found two types of rheological behavior, namely that of a viscoelastic liquid and that of a soft solid. Quite remarkable, these two behaviors are observed on the same samples, typically in a ratio 60%-40%. While a similar elastic modulus was obtained in viscoelastic liquid and soft solid regimes, of the order of 2 - 5 Pa, the relaxation times (the ratio between the viscosity and the modulus) are different, one being measurable at about 40- 50 s, and the other being about an order of magnitude longer (> 300 s). It is concluded that the variability in viscosity that we obtain is related with the spatial distribution of relaxation times in mucus at a scale comprised between 10 and 100 μm (*i.e.* the size of the wires), an outcome that was not previously disclosed and which could be relevant for mucus thinning treatments. We also showed that mucus collection from tissue culture, leading to the late *Ex Vivo* mucus in this work can be successfully used for the investigation of mucus rheology. This suggests that the tissue culture method, but also possibly culture of human lung epithelial cells may be appropriate for investigations related to drug delivery with similar rheological properties to *In Vivo* mucus. This work finally shows that beyond the established structural variations due to the microcavities, mucus exhibits secondary inhomogeneities associated with the relaxation times of the mucin network that may be important for its flow properties.

## Supporting information

SI Radiom et al. submitted to BioRxiv 09-Apr-21

## Acknowledgments

ANR (Agence Nationale de la Recherche) and CGI (Commissariat à l’Investissement d’Avenir) are gratefully acknowledged for their financial support of this work through Labex SEAM (Science and Engineering for Advanced Materials and devices) ANR 11 LABX 086, ANR 11 IDEX 05 02. We acknowledge the ImagoSeine facility (Jacques Monod Institute, Paris, France), and the France BioImaging infrastructure supported by the French National Research Agency (ANR-10-INSB-04, « Investments for the future »). This research was supported in part by the Agence Nationale de la Recherche under the contract ANR-13-BS08-0015 (PANORAMA), ANR-12-CHEX-0011 (PULMONANO), ANR-15-CE18-0024-01 (ICONS), ANR-17-CE09-0017 (AlveolusMimics) and by Solvay.

## Supporting Information

S1: Human mucus samples used in this study – S2: Origin of the human mucus samples – S3: Synthesis of magnetic wires – S4: Characterization of magnetic wires – S5: Effect of pH on the magnetic wire stability – S6: Magnetic rotational spectroscopy: methods: S7 – Magnetic rotational spectroscopy *versus* macrorheology: a comparative study – S8: Analysis of early *Ex Vivo* mucus – S9: Review of previous sputum and mucus rheology

## TOC and graphical abstract

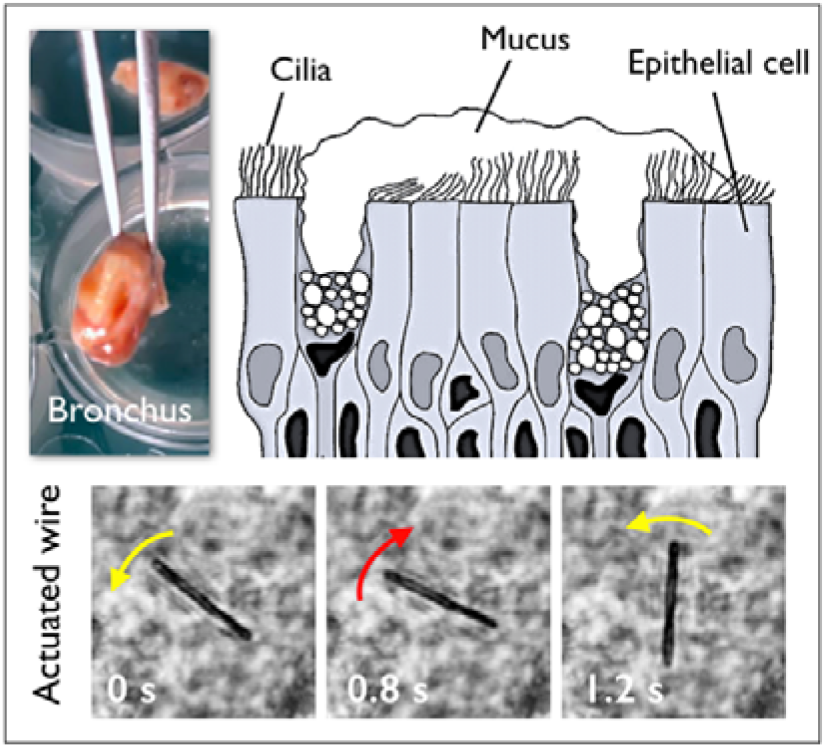

Magnetically actuated micron-sized wires are used to address the question of the mechanical properties of mucus gels collected from human bronchus tubes following surgery. Our work shows that mucus has the property of a high viscosity gel characterized by large spatial viscoelastic heterogeneities.

