## Supplementary material for "Magnetic wire active microrheology of human respiratory mucus": SI Radiom et al. submitted to BioRxiv 09-Apr-21

**Outline**

**SI 1** – Human mucus samples used in this study

**SI 2** – Origin of the human mucus samples

**SI 3** – Synthesis of magnetic wires

**SI 4** – Characterization of magnetic wires

**SI 5** – Effect of pH on the magnetic wire stability

**SI 6** – Magnetic rotational spectroscopy: methods

**SI 7** – Magnetic rotational spectroscopy *versus* macrorheology: a comparative study

**SI 8** – Analysis of early *Ex Vivo* mucus

**SI 9** – Review of previous sputum and mucus rheology

**Movie#1** – Wire rotation associated with Fig. 1a

Movie showing a 23 µm wire undergoing rotation in *Ex Vivo* mucus as a result of a 10 mT external magnetic field rotating with an angular frequency of 0.0094 rad s^-1^.

**Movie#2** – Wire rotation associated with Fig. 1b

Same as movie#1 for an angular frequency of 0.94 rad s^-1^.

**Movie#3** – Wire rotation associated with Fig. 4a

Movie showing a 23 µm wire undergoing rotation in Ex Vivo mucus as a result of a 10 mT external magnetic field rotating with an angular frequency of 0.0031 rad s^-1^.

**Movie#4** – Wire rotation associated with Fig. 4b

Same as Movie#3 at 0.0094 rad s^-1^.

This version Friday, April 9, 2021 - Submitted to BioRxiv

**Supporting Information S1**

Human mucus samples used in this study

| **Sample** | **Collection** | **Conditioning** |
| --- | --- | --- |
| Early *Ex Vivo* | Directly from bronchus tube | Room temperature, washed in Tyrode solution |
| Late *Ex Vivo* | From *Ex Vivo* culture of excised bronchus tube | 12 – 18 h incubation (37°C, 5% CO_2_) |

***Table S1****: Definitions of mucus samples.*

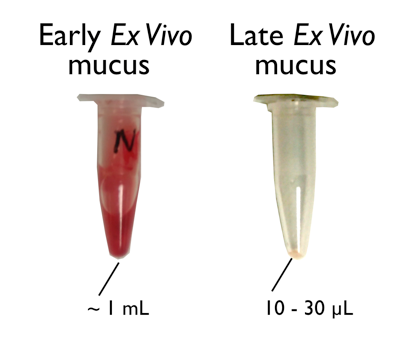

***Figure S1****: Images of Eppendorf containing the early and late Ex Vivo mucus*

**Supporting Information S2**

Origin of the human mucus samples

| **Date of surgery** | **Sex** | **Age** | **Pulmonary diseases** | **Samples** |
| --- | --- | --- | --- | --- |
| 23/01/2019 | M | 61 | Pulmonary adenocarcinoma | Late *Ex Vivo*  Early *Ex Vivo* |
| 28/01/2019 | M | 58 | Pulmonary hypertension and fibrosis | Late *Ex Vivo*  Early *Ex Vivo* |
| 21/03/2019 | F | 56 | Pulmonary fibrosis | Late *Ex Vivo*  Early *Ex Vivo* |
| 16/04/2019 | M | 66 | Pulmonary fibrosis | Late *Ex Vivo*  Early *Ex Vivo* |
| 21/05/2019* | M | n.s. | none | Early *Ex Vivo* |
| 21/05/2019 | M | 56 | Pulmonary fibrosis | Early *Ex Vivo* |
| 06/06/2019 | M | 48 | Pulmonary hypertension and fibrosis | Late *Ex Vivo* |
| 04/07/2019 | M | 62 | Pulmonary hypertension and fibrosis | Late *Ex Vivo*  Early *Ex Vivo* |

***Table S2****: Patients pathologies. This patient designated with a star* was a donor with no identified respiratory disease. n.s. is for “not specified”.*

**Supporting Information S3**

Synthesis of magnetic microwires

Superparamagnetic maghemite ($\gamma$-Fe_2_O_3_) nanoparticles with a mean diameter of 13.2 nm and a dispersity of 0.23 are used (**Fig. S3-1**). The nanoparticles are initially coated with poly(acrylic) acid polymer at pH 2.0. The precipitate is then re-dispersed by increasing the pH to 10. The negatively charged particles are then assembled using poly(diallydimethyl ammonium chloride) (PDADMAC) polycations. The cylindrical shape was given during desalting the mixed solutions using a dialysis cassette under a magnetic field of 300 mT (**Fig. S3-2**).

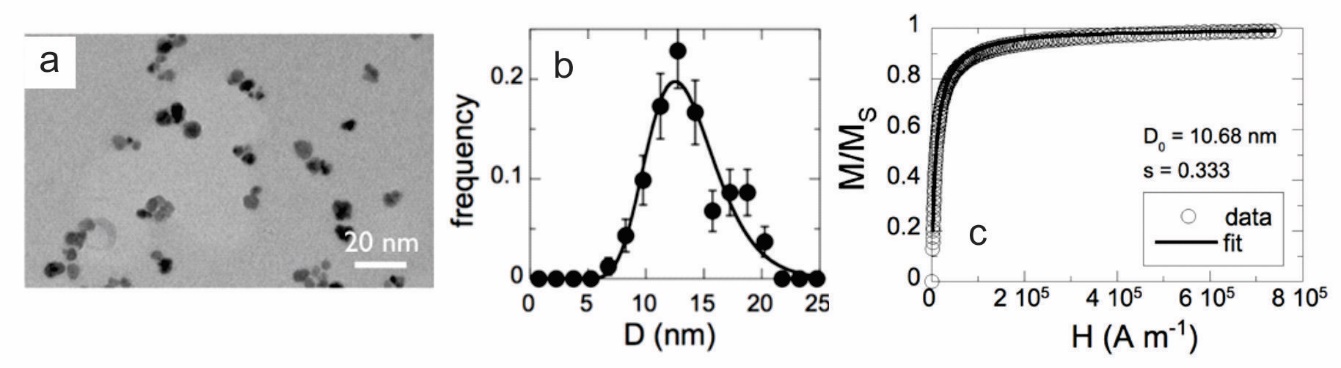

***Figure S3-1****: (a-c) Iron oxide nanoparticles: transmission electron microscopy image (a), size distribution (b), and magnetic field H-dependence of the macroscopic magnetization M(H) normalized by its saturation value M_S_ for cationic (uncoated) maghemite dispersions (c). The experiment was performed using vibrating sample magnetometry (VSM). The solid curve was obtained using the Langevin equation convoluted with a log-normal distribution of particle sizes. M_S_ =ϕ × m_s_, where m_s_ is the specific magnetization of colloidal maghemite (m_s_ = 3.5×10^5^ A m^-1^) and ϕ the volume fraction. The nanoparticle diameter derived from VSM and from TEM are different,* $D_{TEM}$ *= 13.2 nm versus* $D_{VSM}$ *= 10.7 nm (1,2).*

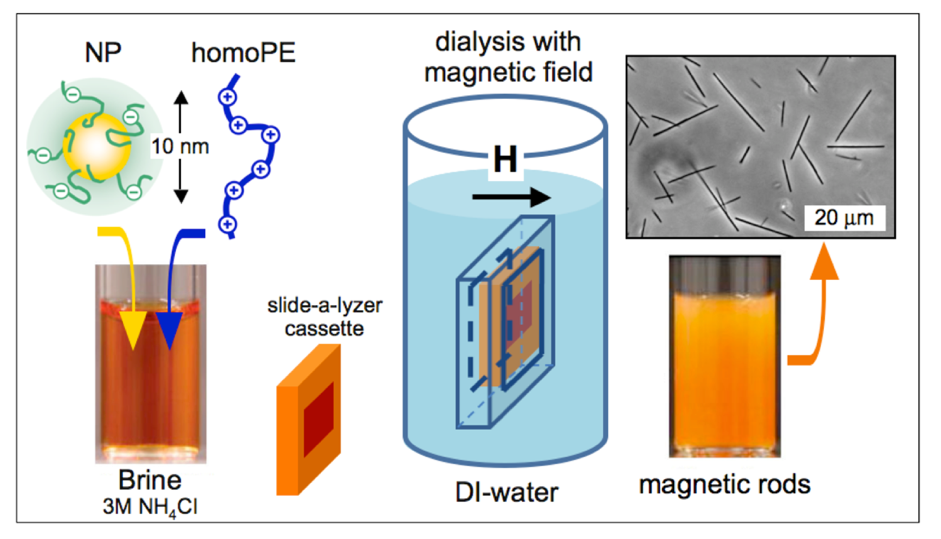

***Figure S3-2****: Schematic representation of the protocol that controls the nanoparticle co-assembly and wire formation. The dialysis involves the preparation of separate 1 M NH_4_Cl salted solutions of particles and oppositely charged polymers. The ionic strength is progressively diminished by dialysis with a 10000 g mol^-1^ cut-off Slide-a-Lyzer cassette. Light microscopy image of the wires is shown.*

**Supporting Information S4**

Characterization of magnetic microwires

The geometrical characterization of the microwires was performed by measuring the length (*L*) and the diameter (*D*) of wires using a 100× objective lens on an optical microscope (Olympus IX73) coupled with a CCD camera (QImaging, EXi Blue) supported by the software Metaview (Universal Imaging). From the distribution, we calculate the reduced wire length, $L^{*}=L/\left[ D\sqrt{g(L/D)} \right]$, where $g\left( x \right)=ln\left( x \right)-0.662+0.917x-0.050x^{2}$ (**Fig. S4-1**).

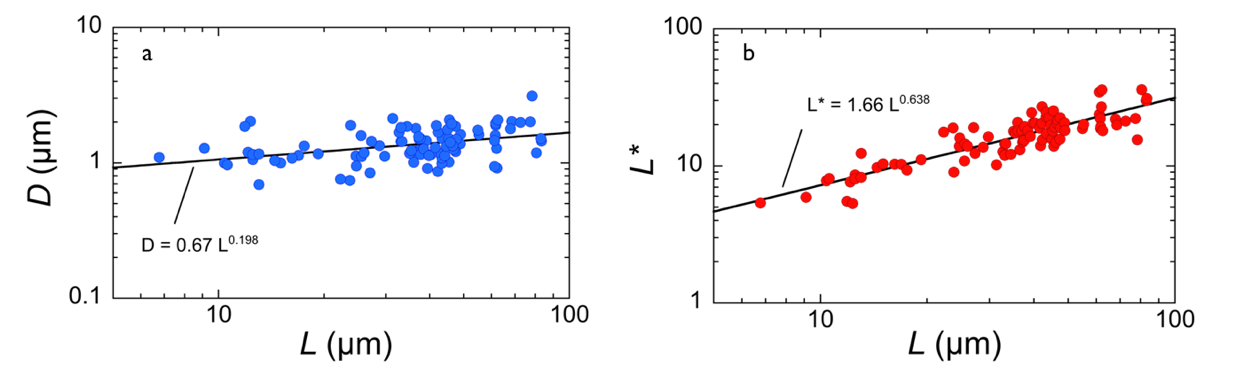

***Figure S4-1****: Distribution of the reduced wire length (*$L^{*}$*) as a function of the wire length (*$L$*). The distribution is fit to a power law giving* $L^{*}=1.66L^{0.64}$*.*

The anisotropy ratio ($\Delta\chi$) of the wires was calculated in independent measurements in fluids of known viscosity, such as water-glycerol mixtures. In this case, by collecting the critical frequency $\omega_{C}$ for wires of various reduced length $L^{*}$, $\Delta\chi$ was calculated using equation (2) in the main text. For these wires $\Delta\chi$ was found to be equal to 2.3 (**Fig. S4-2**).

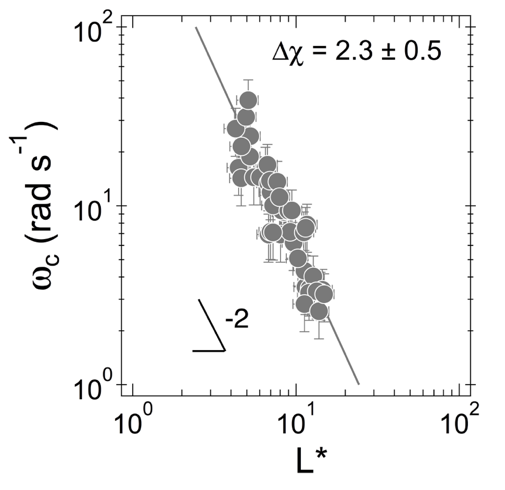

***Figure S4-2****: Distribution of the critical frequency (*$\omega_{C}$*) as a function of the reduced wire length (*$L^{*}$*) for wires in 85% glycerol-water solution of viscosity 0.062 Pa s^-1^ (T = 26 °C). The distribution is fit to Eq. 1 in the main text giving anisotropy ratio* $\Delta\chi$ *= 2.3 ± 0.5 (3).*

**Supporting Information S5**

Effect of pH on the magnetic wire stability

There are indications that airway defenses are affected by the pH of the lung fluids (4). The pH of the airway mucosa has been measured and depending on the pathology the pH was found to vary between 5.5 and 8.3. In this context of microrheology measurements using magnetic wires, it is important to assess the stability of magnetic wires as a function of the pH. The pH of a dispersion containing 15 µm long wires (median value) was modified by addition of hydrochloric acid (pH 1.5, 3.4, 4.1) or sodium hydroxide (pH 9.1). After 4 days, the wires were observed by phase-contrast microscopy and compared to the neutral pH conditions (5-7). **Fig. S5** shows that from pH 9.1 down to pH 3.4 the wires remained intact, and comparable to those of neutral conditions (pH 7.5). At pH 1.4, the wires changed slightly but were still not degraded. Few wires exhibit kinks and bending that indicate a softening of their structure.

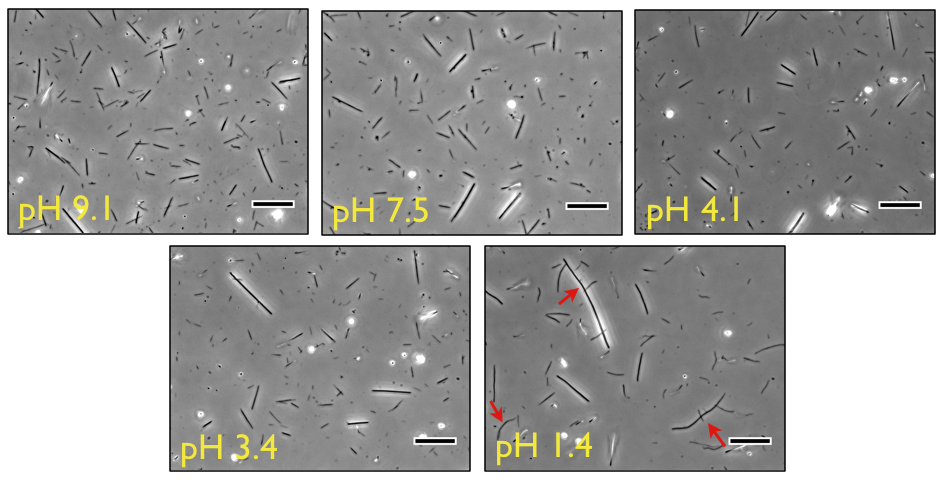

***Figure S5****: Phase-contrast images of magnetic wires at different pHs between pH 9.1 and pH 1.4.*

**Supporting Information S6**

Magnetic rotational spectroscopy methods

The magnetic device consists of two pairs of coils each 23 Ω. The current input to the coils is by a two-channel frequency generator and an amplifier. Using this setup which is schematically shown in **Fig. S6**, we applied rotating magnetic fields with an amplitude of 10.3 mT and a rotational velocity ranging from 0.001 – 10 rad s^-1^ to the wires in mucus. The evolution of wire orientation angles was acquired on IX73 Olympus inverted microscope with 20× objective lens and further analyzed using plug-ins in ImageJ.

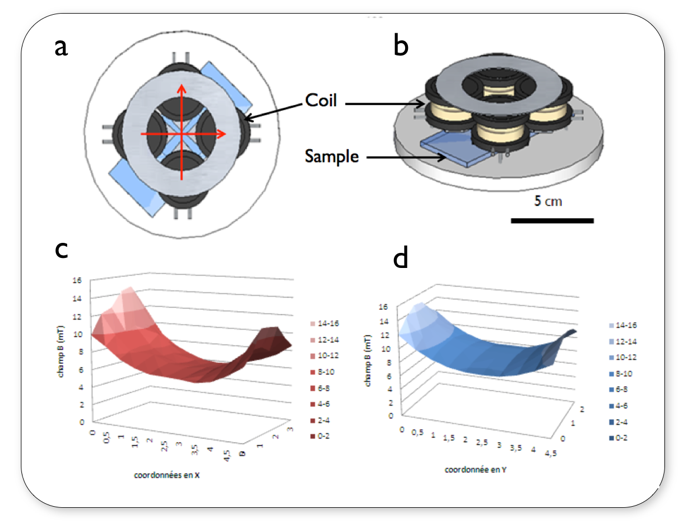

***Figure S6****: Top (a) and side (b) views of the rotating field device used in this work. Magnetic field distributions are shown along the X (c) and Y (d) axis of the four-coil device. In the center, the magnetic field is constant over a 1*$\times$*1 mm^2^ range.*

**Supporting Information S7**

Magnetic rotational spectroscopy *versus* macrorheology: a comparative study

*Microrheology with a viscoelastic liquid model*

A mixture of cetylpyridinium chloride (CPCl) and sodium salicylate (NaSal) dispersed in 0.5 M NaCl solution at a concentration of 2 wt.% forms a perfect (Maxwell) viscoelastic liquid. CPCl and NaSal are known to self-assemble spontaneously into micrometer long wormlike micelles, which then build a semi-dilute entangled network with a mesh size of 30 nm at the used concentration (8,9). **Fig. S7-2a** displays a cone-and-plate device (diameter 50 mm, angle of the cone 2°, CSL 100 rheometer, TA Instruments) used in macrorheology and **Fig. S7-2b** the frequency dependence of the elastic and viscous moduli $G^{'}\left( \text{ω} \right)$ and $G^{''}\left( \text{ω} \right)$ obtained with this solution. At 27 °C, the solution was characterized by a static viscosity $\eta_{0}=1.0\pm0.1$ Pa s, and an elastic modulus $G_{0}=7.1\pm0.1$ Pa (8). The relaxation time of the solution is then estimated to be $\tau_{R}=0.14$ s. The continuous lines are the predictions for a Maxwell fluid: ${G^{'}\left( \text{ω} \right)}/{G_{0}={X^{2}}/\left( 1+X^{2} \right)}$ and ${G^{''}\left( \text{ω} \right)}/{G_{0}=X/\left( 1+X^{2} \right)}$ with $X=\omega\tau_{R}$. The agreement between the data and Maxwell model predictions is excellent. In **Fig. S7-2c and S7-2d**, a rotating magnetic field of 10.4 mT was applied to 8.1 µm wire (inset) immersed in the solution at increasing frequencies 0.1 – 20 rad s^-1^. The motion of the wire was monitored by optical microscopy, and the time dependence of orientation angle derived. **Fig. S7-2c** show 4 experimental time traces $\theta\left( t \right)$ obtained at excitation frequencies 0.14, 0.40, 2.9 and 17.0 rad s^-1^. At low frequency, the wire rotates with the field, and $\theta\left( t \right)=\omega t$. Above the critical frequency (here $\omega_{c}=0.38$ rad s^-1^), the wire performs turn-and-return response behavior characteristic of the asynchronous regime. The red straight lines in the figures represent the average angular velocity $\Omega\left( \omega\right)$. **Fig. S7-2d** displays the evolution of the average rotational velocity versus excitation frequency obtained with several wires investigated in this fluid. The data were adjusted using equation (1) and (2) in the main text and a value of the viscosity equal to 1.3 ± 0.3 Pa s, in good agreement with cone-and-plate rotational rheometry is obtained. The comparison between macro- and microrheology shows that rotating wires are able to account for the static shear viscosity of a Maxwell fluid.

*Microrheology with a viscoelastic solid model*

Phytagel powder was added slowly to 1 mM calcium chloride solution at room temperature with rapid stirring and heated up to 50^°^C. A final concentration equal to 0.3 wt.% was prepared higher than the sol to gel transition concentration. The sample was studied by cone-and-plate rheometry in the same conditions as the surfactant micelles. **Fig. S7-3a** displays the cone-and-plate geometry used for the rheological measurements and **Fig. S7-3b** the frequency dependences of the elastic and viscous moduli. We find that $G^{'}\left( \text{ω} \right)$ and $G^{''}\left( \text{ω} \right)$ exhibit scaling behaviors with exponents 0.20 and 0.15, respectively. In addition, on the whole frequency range, the inequality $G^{'}\left( \text{ω} \right)> G^{''}\left( \omega\right)$ is found. These two properties are known to be indicators of a viscoelastic solid behavior (10,11). In microrheology using MRS, wires of lengths 6 – 45 µm were immersed in the sample and submitted to a rotating magnetic field of 12 mT at a frequency between 5×10^-3^ – 10 rad s^-1^. **Fig. S7-3c** shows time traces of the orientation angle obtained at 0.015, 0.15, 1.54 and 6.83 rad s^-1^. Over 3 decades in frequency, the traces reveal a unique behavior: $\theta\left( t \right)$ displays regular oscillations at a frequency double of that of the field, and an average rotational velocity $\Omega\left( \omega\right)$ (shown as red straight lines). **Fig. S7-3d** displays the average rotational velocity *vs* the excitation frequency, and indicates that for all frequencies tested, $\Omega\left( \omega\right)\cong0$ within the measurement uncertainty. This behavior agrees with the prediction for the Kelvin-Voigt model. From the amplitude of the oscillation $\theta_{B}\left( \omega\right)$, the elastic modulus $G=2.5\pm2.8$ Pa is estimated using the expression of equation (3) in the main text in good agreement with the rheological cone-and-plate value $G^{'}=3.0\pm0.5$ Pa. In conclusion, we show that our MRS technique is able to account for the time and frequency dependencies of a viscoelastic solid of Kelvin-Voigt type.

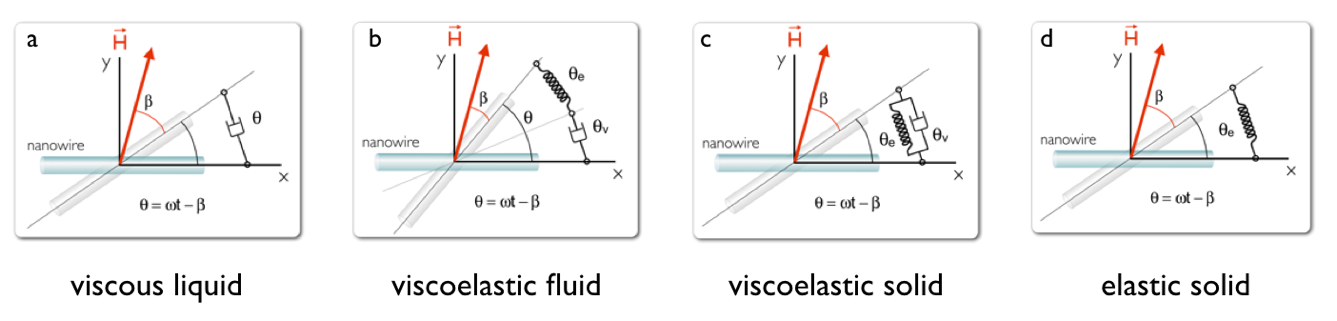

***Figure S7-1****: Schematic representation of a wire in a purely viscous liquid (a), viscoelastic liquid (b), viscoelastic solid (c) and elastic solid (d). The viscous liquid is shown as a dashpot and the elastic solid as a spring. A spring and a dashpot in series form a Maxwell element. A spring and a dashpot in parallel form a Kelvin-Voigt element.*

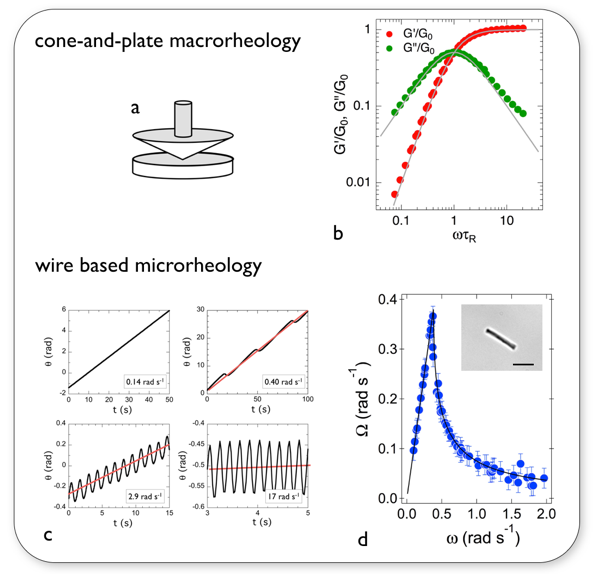

***Figure S7-2****: (a) Cone-and-plate geometry used in shear rheometry. (b)* ${G^{'}\left( \text{ω} \right)}/{G_{0}}$ *and* ${G^{''}\left( \text{ω} \right)}/{G_{0}}$ *vs frequency for a wormlike micellar fluid. The continuous lines are Maxwell predictions. (c) Orientation angle* $\theta\left( t \right)$ *of 8.1 µm wire as a function of the time and at several actuation frequencies 0.14, 0.40, 2.9 and 17.0 rad s^-1^. (d) Average rotational velocity* $\Omega\left( \omega\right)$ *as a function of the frequency. The solid line corresponds to the best fit using equation (1) in the main text. Inset in (d): image of the wire by phase contrast microscopy (60×, scale bar 5 µm).*

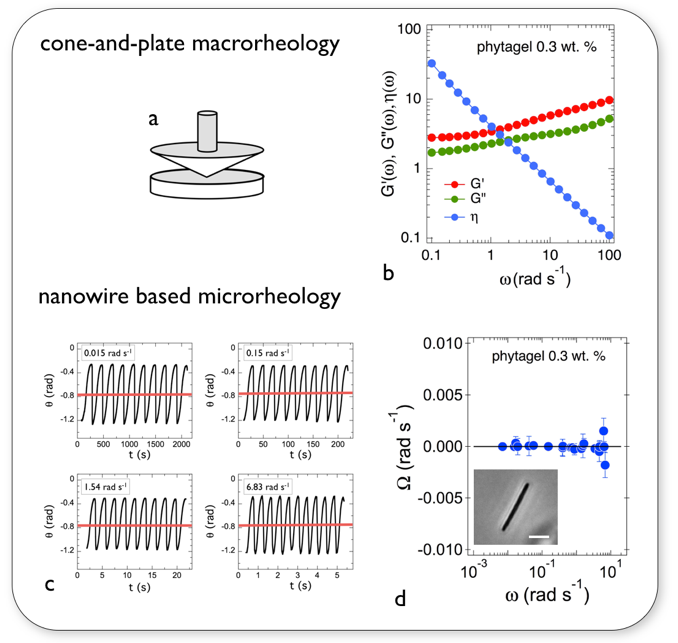

***Figure S7-3****: (a) Cone-and-plate geometry used in shear rheometry. (b)* $G^{'}\left( \text{ω} \right)$*,* $G^{''}\left( \text{ω} \right)$ *and* $\eta\left( \text{ω} \right)$ *vs frequency of a 0.3 wt.% phytagel sample. (c) Orientation angle* $\theta\left( t \right)$ *of 12 µm wire as a function of the time for various values of actuation frequency 0.015, 0.15, 1.54 and 6.83 rad s^-1^. (d) Average rotational velocity* $\Omega\left( \omega\right)$ *as a function of the actuation frequency. The solid line corresponds to the best fit using the Kelvin-Voigt model. Inset in d): image of the 12 µm wire by phase contrast microscopy (60×, scale bar 5 µm).*

**Supporting Information S8**

Analysis of early *Ex Vivo* mucus

Early *Ex Vivo* mucus gels were studied using MRS following the protocol described in the main text as for late Ex Vivo mucus. The proportions of the two generic behaviors are 35 cases (first) to 23 cases (second) in a total 58 wires investigated in 8 different samples. **Fig. S7-1a** shows the average rotational frequency $\Omega\left( \omega\right)$ normalized with the critical frequency as a function of the reduced actuation frequency $\omega/\omega_{C}$. As for late *Ex Vivo* mucus samples, the data are well accounted for by the viscoelastic model prediction (continuous curve). Fig. **S7-1b** displays $\theta_{B}\left( \omega\right)$ as a function of the normalized frequency $\omega/\omega_{C}$. It is found that the data superimpose over 4 decades in frequency and follow a scaling law with exponent -0.12.

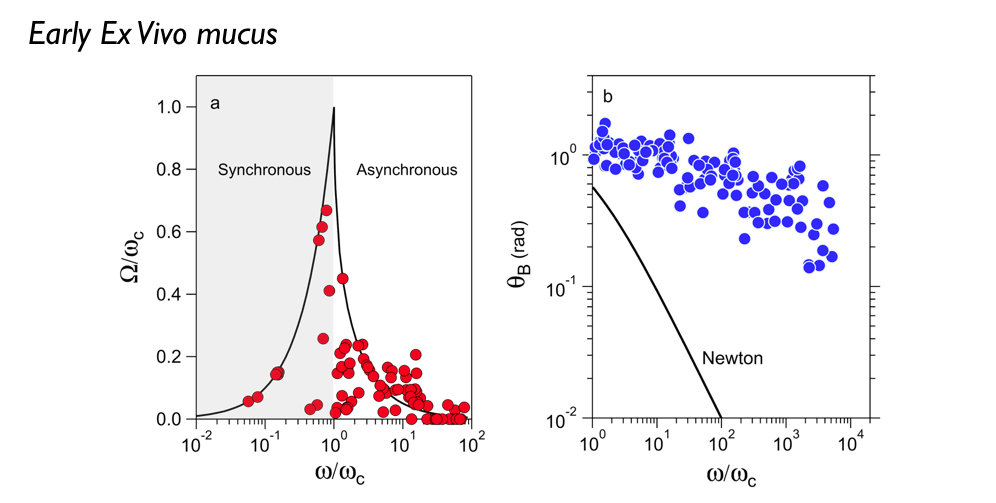

***Figure S8-1****: a) Average rotational velocity of wires normalized by their individual critical frequency* $\Omega\left( \omega\right)/\omega_{C}$ *plotted versus normalized frequency* $\omega/\omega_{C}$*. The response behaviors of the wires (35 wires) collapse on a master curve. b) Oscillation amplitude* $\theta_{B}$ *as a function of normalized frequency* $\omega/\omega_{C}$ *and the difference between response in mucus and response in a Newtonian liquid*

**Fig. S8-2a** displays $\omega_{C}$ as a function of $L^{*}$ for the same wires. In this figure, each data point represents a single wire embedded in a different mucus environment. Using Eq. 1 we calculated the static viscosity of the individual wires and obtained a mean value equal to 180 ± 30 Pa s, again in agreement with the late Ex Vivo data. **Fig. S8-2b** depicts the angle $\theta_{0}$ for 35 individual wires as a function of the reduced wire length and the $1/{L^{*}}^{2}$-dependence of this angle. Using Eq. 2, we calculated the elastic modulus and obtained a mean of 4.4 ± 0.5 Pa. A similar calculation was made for the low frequency region, from which the equilibrium modulus can be derived *via* the relation. It is found that $G_{eq}$ = 2.0 ± 0.2 Pa (**Fig. S8-2c**).

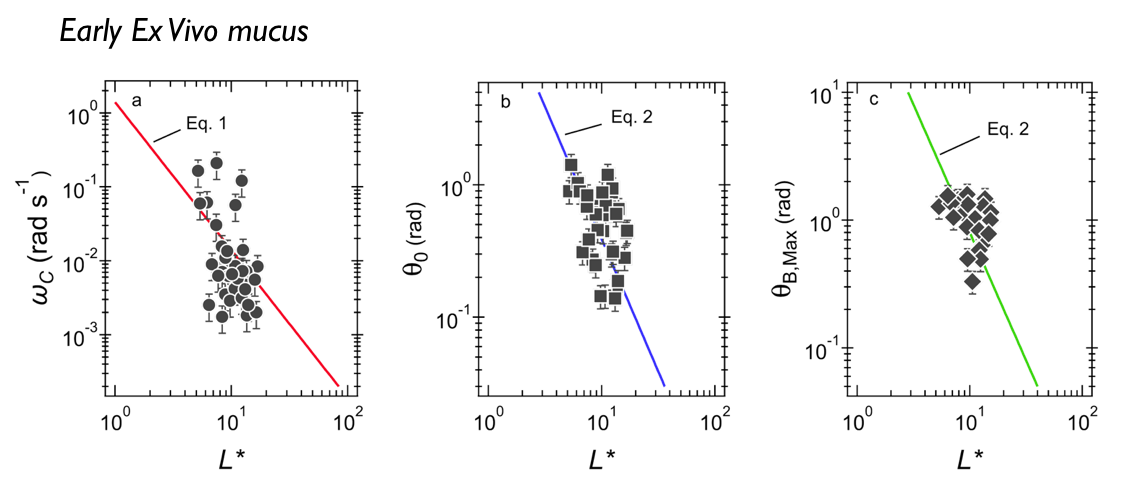

***Figure S8-2****: a) Variation of critical frequency* $\omega_{C}$ *as a function of reduced wire length* $L^{*}$*. The thick solid line shows the* $1/{L^{*}}^{2}$*-dependence corresponding to a mean viscosity* $\eta$*=* 80 ± 30 *Pa s. b) Variation of* $\theta_{0}$ *as a function of reduced wire length* $L^{*}$*. Solid line shows the* $1/{L^{*}}^{2}$*-dependence of Eq. 2, leading to* $G_{0}$ *=* 4.4 ± 0.5 *Pa. c) Variation of oscillation amplitude* $\lim_{\omega\to0} \theta_{B}\left( \omega\right)$ *as a function of the reduced wire length. Solid line shows the* $1/{L^{*}}^{2}$*-dependence of Eq. 2, leading to* $G_{eq}$ *=* 2.0 ± 0.2 *Pa.*

**Supporting Information S9**

Review of previous sputum and mucus rheology

Macro and microrheological data from the literature were evaluated for sputum and mucus from different human and animal sources and from related diseases, such as chronic obstructive pulmonary disease (COPD) and cystic fibrosis (CF). The list is provided in Table S1.

|  | **Mucus** | | | **Sputum** | | |  |
| --- | --- | --- | --- | --- | --- | --- | --- |
| **References** | **Healthy** | **COPD** | **CF** | **Healthy** | **COPD** | **CF** | **Cells** |
| Dulfano-71 (12) |  |  |  |  | human |  |  |
| King-81 (13) |  |  |  |  |  | canine |  |
| Puchelle-81 (14) |  |  |  |  |  | human |  |
| Jeanneret-Grosjean-88 (15) | human |  |  |  |  |  |  |
| Shah-96 (16) |  |  |  |  |  | human |  |
| Griese-97 |  |  |  |  |  | human |  |
| Sanders-00 (17) |  |  |  |  |  | human |  |
| Dawson-03 (18) |  |  |  |  |  | human |  |
| Besseris-07(19) | canine |  |  |  |  |  |  |
| Suk-09 (20) |  |  |  |  |  | human |  |
| Seagrave-12 (21) |  |  |  |  |  |  | human |
| Kirch-12 (22) | horse |  |  |  |  |  |  |
| Schuster-13 (23) | human |  |  |  |  |  |  |
| Tomaiuolo-14 (24) |  |  |  |  |  | human |  |
| Vasquez-14 (25) | horse |  |  |  |  |  |  |
| Hill-14 (26) |  |  |  |  | human | human |  |
| Yuan-15 (27) |  |  |  | human |  | human |  |
| Murgia-16 (28) | porcine |  |  |  |  |  |  |
| Requena-17 (29) | human |  | human |  |  |  |  |
| Ma-18 (30) |  |  |  |  |  | human |  |
| Radtke-18 (31) |  |  |  |  |  | human |  |
| Jory-19 (32) | human | human |  |  |  |  |  |

***Table S9:*** *List of publications on the rheology of sputum and mucus. The references are classified chronologically. In the first column, they are labeled with the name of the first author and with the year of publication. The origin of the sputum and mucus samples investigated is indicated.*
